# Multi-Omics Exploration of ABA Involvement in Identifying Unique Molecular Markers for Single and Combined Stresses in tomato plants

**DOI:** 10.1101/2024.05.13.593826

**Authors:** Miriam Pardo-Hernández, Pascual García-Pérez, Luigi Lucini, Rosa M Rivero

**Affiliations:** Center of Edaphology and Applied Biology of Segura (CEBAS-CSIC), Department of Plant Nutrition, Campus Universitario Espinardo, Ed 25 30100, Murcia (Spain); Department for Sustainable Food Process, Università Cattolica del Sacro Cuore, Via Emilia Parmense 84, 29122 Piacenza (Italy)

**Keywords:** multi-omics, salinity, heat, stress combination, ABA-deficient mutant, tomato

## Abstract

Over the past decade, our research group has found that plant responses to combined abiotic stresses are unique and cannot be inferred from studying plants exposed to individual stresses. Adaptive mechanisms involve changes in gene expression, ion regulation, hormonal balance, and metabolite biosynthesis or degradation. Understanding how these mechanisms integrate from stress perception to biochemical and physiological adjustments is a major challenge in abiotic stress signaling studies. Today, vast amounts of -omics data (genomics, transcriptomics, proteomics, metabolomics, phenomics) are readily available. Additonally, each –omic level is regulated and influenced by the others, highlighting the complexity of plant metabolism’s response to stress. Considering abscisic acid (ABA) as a key regulator in plant abiotic stress responses, in our study, ABA-deficient plants (*flc*) exposed to single or combined salinity and heat stresses were evaluated and different -omics analyses were conducted. Significant changes in biomass, photosynthesis, ions, transcripts, and metabolites occurred in mutant plants under single or combined stresses. Exogenous ABA application in *flc* mutants did not fully recover plant phenotypes or metabolic levels but induced cellular reprogramming with changes in specific markers. Multi-omics analysis aimed to identify ABA-dependent, ABA-independent, or stress-dependent markers in plant responses to single or combined stresses. We demonstrated that studying different -omics together identifies specific markers for each stress condition not detectable individually. Our findings provide insight into specific metabolic markers in plant responses to single and combined stresses, highlighting specific regulation of metabolic pathways, ion absorption, and physiological responses crucial for plant tolerance to climate change.

**Highlight:** The integration of different -omics has enabled the identification of specific ABA-dependent or –independent markers for single or combined abiotic stresses. These markers were not initially detectable when studying the different –omics individually.

## Introduction

Tomato (*Solanum lycopersicum* L.) is one of the most consumed vegetable crop globally, valued for its nutritional and health benefits, as well as for being economic significance (Delian *et al*., 2017; Liu *et al*., 2022). Widely considered a primary model organism, the tomato plant boasts a small genome size and the availability of hundreds of sequenced genomes, facilitating experimental manipulation (Sun *et al*., 2020; Xia *et al*., 2021). However, its growth and productivity are susceptible to abiotic stresses, including temperature fluctuations, drought, salinity, water logging and heavy metal soil pollution (Naik *et al*., 2023). Throughout their whole life cycle, plants have to face different abiotic and biotic stresses, which often act simultaneously rather than independently (Husaini, 2022; López-Valdez *et al*., 2022). Studies over the last decade have shown that plant responses to the combination of two or more abiotic stresses are unique and cannot be inferred from the study of plants exposed to each stress separately (Zandalinas *et al*., 2021; Pascual *et al*., 2022; Rivero *et al*., 2022; Pardo-Hernández *et al*., 2024). In addition, climate change and global warming are influencing crop yield via exacerbated biotic and abiotic stresses, and therefore, many scientists are investigating how to mitigate climate change effects or enhance the stress tolerance of plants (Shahzad *et al*., 2022; Asif *et al*., 2023). Plants respond to these stresses through adaptive mechanisms that involve changes in gene expression, ion regulation, hormonal balance, proteins and metabolite biosynthesis or degradation, among others. The alterations observed under a given abiotic stress or under stress combination in each of these metabolic processes are specific to each stress, as well as to its intensity and duration (Gong *et al*., 2020; Jiang *et al*., 2022). Today, vast amounts of -omics data (genomics, transcriptomics, proteomics, metabolomics, phenomics) are readily available, offering insights into plant stress responses. Understanding how these mechanisms integrate from stress perception to biochemical and physiological adjustments is a major challenge in abiotic stress signaling studies. Additonally, each – omic level influences and is influenced by others, highlighting the complexity of plant metabolism’s response to stress. Thus, multi-omics has developed as a new tool to understand the complexity of the regulation of this intricate metabolic network, and aims to identify specific molecular markers in each –omic level by the existing and known connections among of all of them (Subramanian *et al*., 2020; Derbyshire *et al*., 2022; Roychowdhury *et al*., 2023).

An important regulator to abiotic stress response in plants is the phytohormone abscisic acid (ABA), which is involved in numerous metabolic processes, such as stomatal closure, plant development and stress-related genes expression (Fujii *et al*., 2007; Dittrich *et al*., 2019). To understand the ABA-dependent response, mutants deficient in enzymes related to ABA biosynthesis from β-carotene have been generated in several plant species (Brookbank *et al*., 2021). Among them, *flacca* (*flc*) in tomato plants, and *aba3* in *Arabidopsis*, are characterized by mutations in the molybdenum cofactor (MoCo) sulfurase gene (Sagi *et al*., 1999, 2002; Min *et al*., 2000), which involve the final oxidation step of abscisic aldehyde to ABA. It is well known that endogenous ABA concentration increases when plants are subjected to different abiotic stresses, with drought stress being the most studied (Wilkinson and Davies, 2002). Thus, the use of these mutants has enabled the identification of many components of the ABA-dependent pathway under any given stress conditions. This ABA-dependent pathway is involved in many processes, such as growth enhancement (Cutler *et al*., 2010), increased reactive oxygen species and anthocyanins (Watkins *et al*., 2017; Watanabe *et al*., 2018), decreased Ca^2+^ levels (Khan *et al*., 2013), increase transcription of oxidative stress-related genes (Liu *et al*., 2019b) or increased soluble sugars (Xu *et al*., 2023), among others. Nevertheless, plants can also respond to stresses in an ABA-independent way. Our research group recently demonstrated that under abiotic stress combination and ABA-deficiency, an ABA-independent regulation of certain genes encoding for transcription factors or proteins involved in the stress response of plants occurs (Pardo-Hernández *et al*., 2024). For instance, other researchers have showed that ubiquitin ligases ATL31 and ATL6 in *Arabidopsis* and choline monoooxygenase in *Kochia scoparia* regulate plant response to salt stress through an ABA-independent pathway (Kalinina *et al*., 2012; Du *et al*., 2023). Therefore, not only research based on ABA-dependent but also on ABA-independent plant responses is essential to engineer plants with increased tolerance to extreme abiotic stresses due to climate change.

The present study was designed to investigate the involvement of ABA in the multi-omics integrated response (phenomic, ionomic, transcriptomic and metabolomics) of tomato plants to single or combined stress conditions, using tomato ABA-deficient mutants (*flacca*, *flc*, SOLYC07G066480). Exogenous ABA was also added to *flc* mutant plants to confirm if ABA was involved in the changes observed in the different -omics analysed. This study revealed that there is a specific response of the tomato plants depending on the stress applied and the endogenous ABA cellular levels. Moreover, the response of ABA-deficient plants under stress combination was not the sum of these effects resulting from single stresses, but a specific response. In this line, *flc* mutant plants treated with exogenous ABA did not fully recover to wild-type plant phenotype, showing an independent response at all omics levels. More importantly, our study revealed that, considering just one -omic level, the molecular markers found might be different from those obtained when considering all the –omics layers as a whole. Thus, our study identified some important molecular markers under control and every stress condition applied, which might serve as fast-track tool for scientists working in the field of abiotic stress and/or ABA.

## Material and methods

### Plant growth and stress treatments

The experiments were carried out on wild-type (WT) and the ABA-deficient mutants *flacca* (*flc*) (donated by TGRC, UC Davis, CA, USA, accession LA4479) of *Solanum lycopersicum* plants (cv Micro-Tom). *Flc* mutants are deficient in the synthesis of the molybdenum-cofactor (MoCo) sulfurylase gene (SOLYC07G066480), therefore they have very low levels of endogenous ABA. Plants were sown, germinated, transplanted and grown as described previously by Pardo-Hernandez et al. (2024). Once all plants had four true leaves, half of the plants for each genotype (24 WT and 48 *flc*) were relocated from Chamber A (optimal conditions) to Chamber B. The only difference between the two chambers was the environmental temperature, being in chamber A 25°C and in chamber B 35°C. Simultaneously, half of the plants from each genotype (24 WT and 48 *flc*) were exposed to a nutrient solution supplemented with 100 mM NaCl. Thus, the following treatments were applied to all plant groups: control (C, 0 mM NaCl and 25 °C), salinity (S, 100 mM NaCl and 25 °C), heat (H, 0 mM NaCl and 35 °C), and salinity + heat (S + H, 100 mM NaCl and 35 °C). In parallel, half of our *flc* mutants of each treatment (24 plants, 6 plants per treatment) received an exogenous application of 100 µM ABA. Twelve days after the stress was applied, the plants were divided into roots and leaves, and the fresh weight (FW) was recorded. Half of the leaves were frozen at -80°C for transcriptomic and metabolomics analysis and the other half were placed in a 60°C oven for ionomic analysis.

### ABA quantification

Endogenous ABA concentration was quantified as described previously by Pardo-Hernández et al. (2024).

### Photosynthetic parameters

The photosynthetic parameters were measured at the end of the experiment in the youngest fully expanded leaf of each plant (three plants per treatment as described previously by Martinez et al. (2018).

### Ionomic analysis

100 mg of dried leaves from three biological replicates were digested with HNO_3_:H_2_O_2_ (5:3; v,v) for cation extraction and another 100 mg from same samples were extracted with water for anion extraction. Anions and cations were determined as described previously by Lopez-Delacalle et al. (2020).

### Transcriptomic analysis

Total RNA of three biological replicates for each treatment and genotype was extracted from 100 mg of frozen tomato leaves using the NucleoSpin^®^ RNA kit (Reference no. 740949.50). Total RNA was quantified, and the RNA integrity number (RIN) was assessed using a bioanalyzer (2100 Expert Plant RNA Nano_DE13806178). For RNA sequencing analysis, 2 µg of total RNA with RIN ≥ 8 from each genotype and treatment were sent to Macrogen (Seoul) for RNA sequencing. The detailed RNA sequencing and subsequent analyses performed were previously described by Pardo-Hernandez et al. (2024). Venn diagrams were created in “jvenn” (Bardou *et al*., 2014).

### Metabolomic analysis

The untargeted metabolomics analysis was analyzed by ultra-high performance liquid chromatography-quadrupole time-of-flight mass spectrometry (UHPLC/QTOF-MS). In short, 100 mg ground and freeze-dried tomato leaf samples were mechanically homogenized in 4 mL of 80% methanol (v/v) acidified with 0.1% formic acid (v/v). Then, extracts were centrifuged at 10,000 g for 10 minutes at 4 °C and the supernatant was filtered by 0.22 µm cellulose filters. For every sample, two technical replicates were examined, making a total of six replicates per experimental group. Plant extracts were analysed by the 6560 Ion Mobility LC/QTOF-MS mass spectrometer (Agilent^Ⓡ^, Santa Clara, CA, USA), setting an injection volume of 6 µL. The operational conditions and the post-acquisition processing data were previously optimized and are described in Supplementary File S1. The database PlantCyc 15.1 by Plant Metabolic Network (pmn.plantcyc.org) was employed for the annotation of reported chemical features, complying with the Level 2 (putatively annotated compounds) of the COSMOS Metabolomics Standards Initiative (Salek *et al*., 2015).

### Multi-omics analysis

MixOmics-derived framework called Data Integration Analysis for Biomarker discovery using Latent variable approaches for Omics studies (DIABLO) (http://mixomics.org/mixdiablo/) (Rohart *et al*., 2017) was used for the integration of four datasets: phenomics, ionomics, transcriptomics and metabolomics, using the software R (v. 4.2.1.). This algorithm confers a supervised approach to identify discriminant features that enable differentiation between treatment groups jointly considering the variability of the four datasets to build three different models (WT, *flc*- and *flc*+). Due to the transcriptomics and metabolomics datasets being very large, differentially expressed genes (DEGs) and differentially accumulated metabolites (DAMs) compared to WT under control conditions were selected for multi-omics analysis. The training parameters were previously optimized as explained earlier (Tomas *et al*., 2024).

### Statistical analysis

The statistical analysis of the experiments was conducted considering, at least, three biological replicates, using the IBM SPSS 26 Statistics software. A one-way analysis of variance (ANOVA) with a p-value < 0.05 was performed for fresh weight, photosynthetic parameters, ABA concentration and ion concentrations, followed by Tukey’s post hoc test. Standard error (SE) values for the different treatments and genotypes were calculated and added in Fig. 1 and Supplementary Table S1. An unsupervised multivariate hierarchical cluster analysis (HCA) was performed to evaluate the similarities and dissimilarities for all treatments based on their transcriptomic and metabolomic profile, separately (using Euclidean distance and Ward’s linkage rule). Significant changes in transcripts and metabolites as compared to the control were defined as log_2_ FC ≥ 121 and 0.05 p-value, according to Volcano analysis to provide DEGs and DAMs (by negative binomial Wald test followed by Benjamini–Hochberg correction in transcriptomic analysis; and by Bonferroni multiple testing correction in metabolomics analysis) (Ferreira and Zwinderman, 2006). As well, a supervised analysis of variance (ANOVA) multi-block orthogonal partial least squares (AMOPLS) was carried out on the metabolomics dataset, using the package rAMOPLS on R (v 4.2.1.) (Boccard *et al*., 2019). The statistical significance of AMOPLS models was set at α = 0.01 and the model was built by performing 100 permutations. The results were expressed as the Relative Sum of Squares (RSS), which represents the percentage of variability ascribed to each factor; Residual Structure Ratio (RSR), which represents the ANOVA consistency of each effect with respect to residuals; RSR p-value, to assess statistical significance; and block contribution, in percentage, associated with each effect. Then, the Variable Importance in Projection (VIP) analysis was used to select the discriminant metabolites attributed to the discrimination due to treatment, plant group and plant group x treatment.

**Fig. 1.**
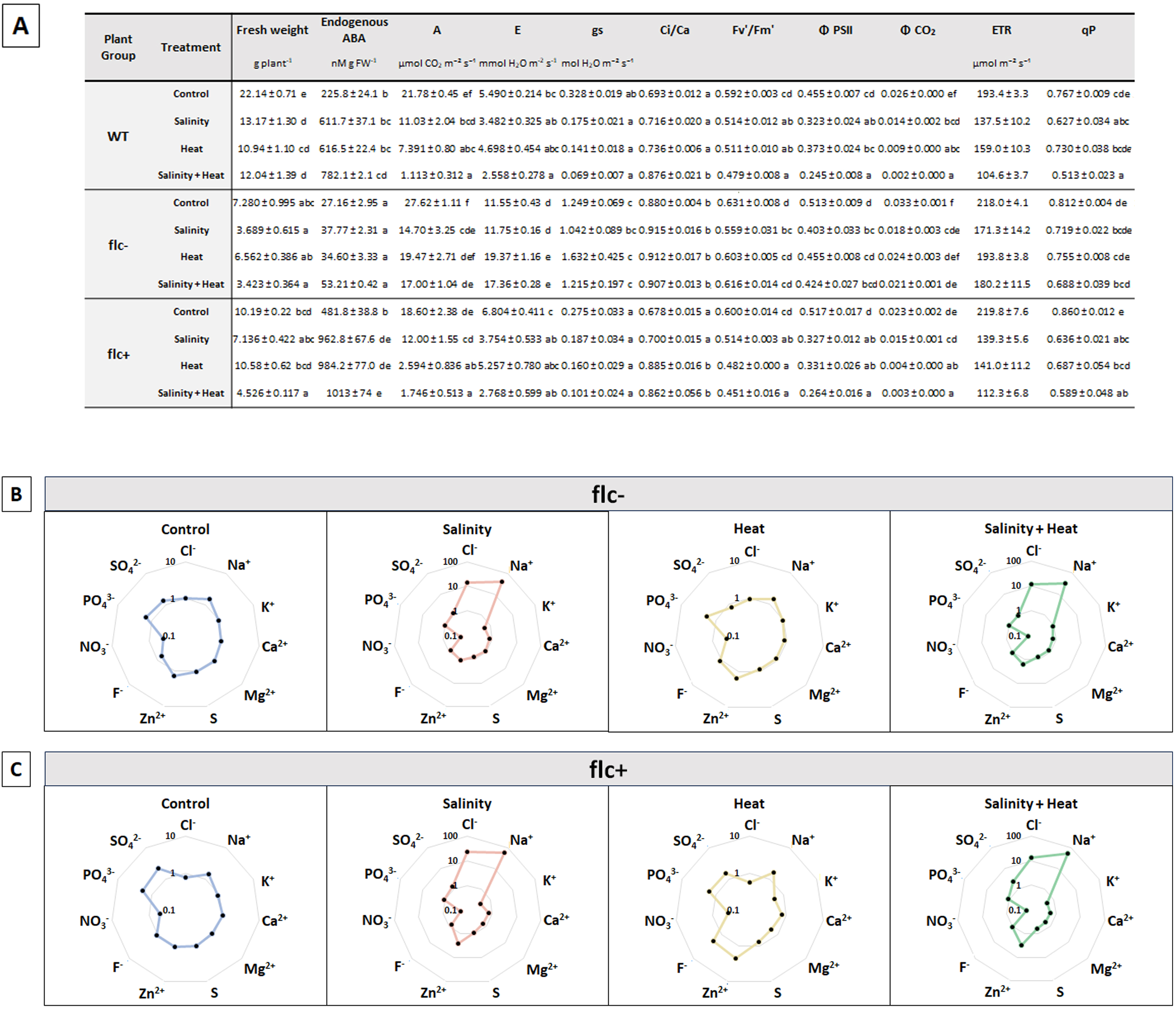
Phenomic and ionomic analysis in tomato wild-type (WT) and ABA-deficient *flacca* mutant without (*flc*-) and with (*flc*+) ABA exogenous application under control, salinity, heat, and combination of salinity and heat. (A) Data of physiological parameters in WT, *flc*- and *flc*+ mutants under control, salinity, heat, and salinity+heat. (B, C) Radial graphs of cations and anions of *flc*- (B) and *flc*+ (C) under control, salinity, heat, and combination of salinity and heat normalized against WT plants grown under control conditions. Raw data for Fig. 1B and 1C can be found in supplemental Supplementary Table S1. A: photosynthesis; E: transpiration; gs: stomata conductance; Ci/Ca: intercellular CO2/ambiente CO2; Fv’/Fm’: Variable to maximal fluorescence (light); Φ PSII: photosystem II quantum yield; Φ CO2: quantum efficiency of CO2 assimilation; ETR: electron transport rate; qP: photochemical quenching.

## Results and discussion

Our research group has been demonstrating for more than a decade that the combination of abiotic stresses produces a specific response in plants that cannot be deduced from the application of the stresses separately. In addition, we have recently demonstrated that, although ABA is an essential hormone in these stress responses, there are certain genetic markers that are regulated independently of ABA and some of them are specific to the combination of salinity and heat (Pardo-Hernández *et al*., 2024). Thus, this work was designed to further investigate the relationship between ABA and the possible coordination of the stress responses of tomato plants at different levels: phenomics, ionomics, transcriptomics, and metabolomics. For this purpose, ABA-deficient tomato mutants (*flc*-) grown under single or combined stress conditions were compared to wild-type (WT) plants. Additonally, to corroborate whether ABA was related to the studied responses at the different –omics levels, half of the *flc* mutants were supplemented with 100 µM ABA and compared to WT plants (*flc*+). Thus, our study aimed to identify different molecular markers specific to each environmental condition in these mutant plants, named as *flc*- and *flc*+ models. Additionally, with the aim of confirming that the identified molecular markers are consistent with the genotype analysed, a WT model was also generated and compared with the previous *flc*- and *flc*+ models generated.

### Tomato response to ABA deficiency under single or combined abiotic stresses: flc- model

#### Phenomics in *flc*-

A phenomic analysis, defined as the physical and biochemical traits defining a genotype (Jin, 2021) was performed to investigate how ABA deficiency might affect the tomato plants under control, salinity, heat and the combination of salinity+heat. Fresh weight, various photosynthetic and chlorophyll fluorescence parameters, and ABA endogenous concentration in tomato leaves were measured on wild-type (WT) and ABA-deficient plants (*flc*-) (**Fig. 1A 1;** Supplementary **Table S1)**.

ABA deficiency in our *flc*- mutants induced a significant reduction in leaf biomass compared to WT plants, which was directly related to inhibition of stomatal closure control and high transpiration rates (Dittrich *et al*., 2019). Controversially, heat stress and the combination of salinity and heat resulted in highest biomass reduction in *flc* mutants respect their control, whereas in WT, single or stress combination resulted in a similar biomass reduction compared to control plants (**Fig. 1A**). As expected, endogenous ABA concentration was significantly lower in *flc*- plants as compared to WT in all the treatments (**Fig. 1A**). Photosynthetic parameters such as CO_2_ assimilation rate (*A*), transpiration rate (*E*), stomatal conductance (*gs*) and intercellular to ambient CO_2_ ratio (*Ci/Ca*) are physiological markers highly correlated with plant growth (Peláez-Vico *et al*., 2024). In our experiments, *A* decreased with all stress treatments in all genotypes respect their controls. *E* and *gs* were significantly increased by ABA deficiency, with the highest values in *flc*-. *Ci/Ca* ratio was also significantly increased in *flc*- plants under control and single stress conditions, but there were no significant differences between the WT and *flc*- plants under stress combination. It has been widely published that transpiration (*E*) is increased in plants stressed by heat to reduce leaf temperature, and reduced in salinity-treated plants (due to associated osmotic stress) (Malini *et al*., 2023), which it was observed in WT plants (**Fig. 1A**). However, and unlike WT, the lack of stomatal control in our *flc*- mutants (Dittrich *et al*., 2019) caused high transpiration rates in salinity-treated plants at the same levels than those observed under control conditions, and even at higher levels under heat stress and under the combination of salinity and heat. Addtionally, chlorophyll fluorescence parameters, such as Fv’/Fm’, PS2 and CO_2_ are also good markers for the stress damage to the photosynthetic machinery. All these parameters were reduced under single or stress combination in WT as compared to control conditions, whereas in *flc*-, only a slightly reduction was observed under salinity stress (**Fig. 1A**). Rock and Zeevaart (1991) working with *aba1-1 Arabidopsis* plants (an homolog of *flacca*), showed that the absence of differences in chlorophyll fluorescence parameters in these mutants could be due to carotenoids, which play a functional or regulatory role in PSII activity (Rock and Zeevaart, 1991). Thus, these pieces of evidence might confirm the results obtained in *flc-* and could prove to be an important tool in elucidating the complex processes that regulate photosynthesis, multi-subunit assembly and energy transfer.

#### Ionomics in *flc*-

A comprehensive ionomics analysis of tomato leaves was performed in WT and *flc*- plants growing under control, salinity, heat and salinity+heat **(Fig. 1B**; Supplementary **Table S1)**. As expected, Cl^-^ and Na^+^ concentrations were significantly higher in salinity and salinity+heat treatments with no differences among the genotypes. K^+^, as an antagonistic ion to Na^+^, exhibited different accumulation patterns depending on whether the stress applied was single or in combination and depending on the genotype studied. In *flc*- plants, K^+^ decreased only under salinity conditions, but not under the combination of salinity+heat. Rivero et al. (2014) showed similar results in a commercial tomato plants, concluding that the specific accumulation of glycine betaine in plants growing under salinity+heat combination prevented Na^+^-induced K^+^ efflux observed under salinity (Rivero *et al*., 2014). Interestingly, Ca^2+^ and Mg^2+^ concentrations followed the same accumulation pattern as described for K^+^ under salinity and salinity+heat conditions. Some authors previously indicated that an application of NaCl might induce a drop in these three cations (K^+^, Ca^2+^, Mg^2+^) (Tzortzakis, 2010; Niu *et al*., 2022) due to their antagonistic relationship within the plant cell. It is worth noting that the NO_3_^-^ concentration was significantly lower in *flc*- under control compared to WT control. ABA has also been established to influence the expression of nitrate reductase (NR), leading to altered NO_3_^-^ influx (Palmer, 1981). Sue et al., (2020) demonstrated that ABA signaling is extensively involved in the regulation of the N starvation response (Su *et al*., 2021). Thus, it is logical to think that our *flc*- mutants are unable to regulate N level within the plant due to ABA deficiency. Pi (inorganic phosphate) contents followed the same pattern as described for Cl^-^ in *flc*- plants; however, in WT plants Pi concentration did not significantly change under any of the stress conditions applied. Unexpectedly, SO_4_^2-^ leaf concentration was only significantly reduced under the combination of salinity+heat in *flc*-, which could be considered as a specific effect of ABA deficiency under stress combination. In short, our results (**Fig. 1B)** demonstrated that, depending on the endogenous concentration of ABA and the stress treatment to which the plant is subjected, concentration of Ca^2+^, K^+^, Mg^2+^, Na^+^, Cl^-^, NO_3_^-^, Pi and SO_4_ ^2-^ differed significantly.

#### Transcriptomics in *flc*-

Metabolically-active leaves were processed for RNA extraction, and an exhaustive RNA sequencing followed by a GO pathway enrichment was performed (Supplementary **Fig. S1**). *Flc*- transcriptome from plants grown under control, salinity, heat and salinity+heat was compared to WT control, obtaining a total of 2578, 3588, 5788 and 4920 differentially expressed genes (DEGs), respectively (**Fig. 2A and** Supplementary **Table S2**). Among these, 344, 610, 1514 and 870 were specifically regulated under control, salinity, heat and stress combination, respectively. In addition, *flc*- plants grown under single stresses, stress combination and control conditions shared 1078 DEGs, indicating genotype-dependent rather than stress-dependent DEGs. Our results also revealed that 161 DEGs were shared by single stresses, 272 DEGs by salinity and stress combination and 1418 DEGs by heat and stress combination (**Fig. 2A**). ABA deficiency led to substantial changes at the transcriptomic level compared to WT in control conditions, with even greater changes observed when *flc*- were subjected to stress conditions, resulting in double the number of DEGs under heat stress and stress combination (**Fig. 2A**).

**Fig. 2.**
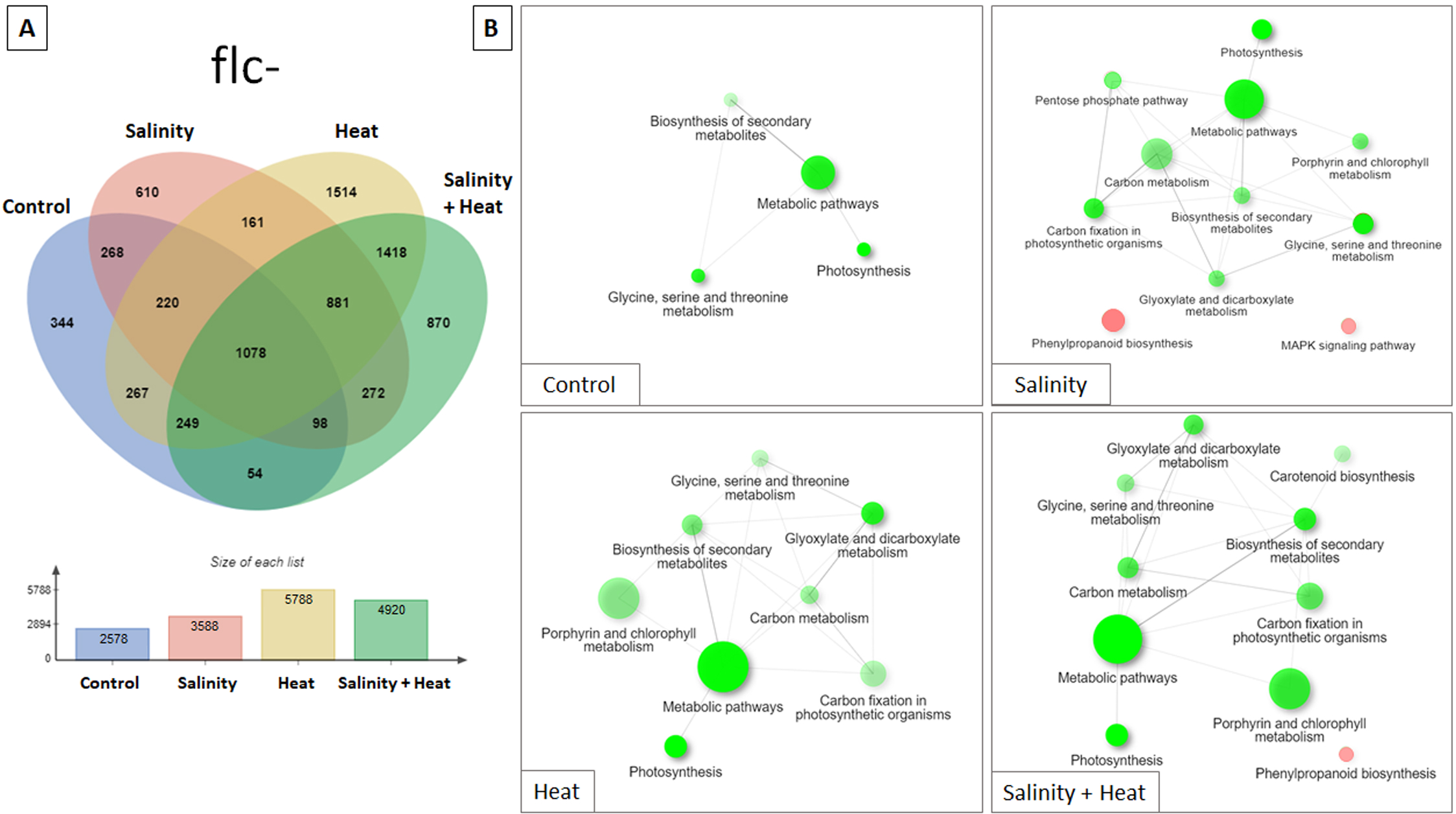
Differentially expressed genes (DEGs) in *flc* mutants under control, salinity, heat and the combination of salinity+heat. DEGs were extracted by normalization against WT control (p<0.05; FC>2). (A) Venn diagrams of the overlap between DEGs in each stress treatment. (B) KEGG pathway enrichment analysis specific DEGs found under control, salinity, heat and salinity+heat treatments in *flc*- mutant plants. The size of the circles is proportional to the number of genes related to that pathway and the color intensity is related to the pathway *P*-value significance level (*P* < 0.05), with red representing upregulated DEGs and green representing downregulated DEGs.

KEGG enrichment pathway analysis (**Fig. 2B**) showed that an ABA deficiency induced a downregulation of genes related to secondary metabolism biosynthesis, photosynthesis, glycine, serine and threonine metabolisms under control conditions. Under single stress or stress combination, *flc*- mutants exhibited significant downregulation of porphyrin and chlorophyll metabolism, carbon metabolism, carbon fixation and glyoxylate and dicarboxylate metabolism was observed. This indicated that transcriptional regulation of these genes was associated with the applied stress conditions (Williams *et al*., 2010; Ashraf and Harris, 2013). Controversely, *flc*- plants grown under salinity showed specific downregulation of the pentose phosphate pathway and upregulation of the phenylpropanoid biosynthesis and MAPK signaling pathways. In contrast, WT plants grown under salt stress exhibited enrichment of the pentose phosphate pathway and MAPK signaling (Miransari *et al*., 2013; Guo *et al*., 2021; Hong *et al*., 2022). Similarly, *flc*- mutants under stress combination displayed specific downregulation of carotenoid biosynthesis and upregulation of phenylpropanoid pathway. Martinez et al (2018) showed that under salinity+heat stress conditions, carotenoid levels and the expression of phenylpropanoid-related genes follow the same trend as observed in our *flc*- plants under these environmental conditions, suggesting an ABA-independent regulation of this pathway (Martinez *et al*., 2018).

#### Metabolomics in *flc*-

To enhance our understanding on how an ABA deficiency might affect plant metabolism under single stresses or stress combination, we conducted an untargeted metabolomics analysis aimed at correlating gene expression data with plant metabolism. This analysis allowed us to positively annotate more than 1515 compounds among WT, *flc*- and *flc*+ (Supplementary **Table S3**). A fold-change-based heatmap from unsupervised hierarchy clustering analysis (HCA) (Supplementary **Fig. S2**) reflected clear differences between plant groups (WT, *flc*-, *flc*+ plants) and treatments (control, salinity, heat and salinity+heat conditions). An ANOVA multi-blocking orthogonal projection to latent structures discriminant analysis (AMOPLS-DA) was conducted to assess which of the three factors (plant group, treatment and plant group x treatment) were the most important in explaining the observed differences (Supplementary **Table S4;** Supplementary **Fig. S3**). Our results indicated that the treatment was the most crucial factor in reflecting differences at the metabolomic level (RSS 21%). Additionally, plant group and the interaction of both plant group and treatment were statistically significant, with a RSS of 16% and 15% respectively. Furthermore, nearly 48% of the observed metabolomic differences could not be explained by these three factors (**Residuals,** Supplementary **Table S4**). Based on AMOPLS-DA, a Variable Important in the Projection (VIP) approach was employed to identify the markers mostly involved in such discrimination (VIP score > 1.2) for each factor. Compounds with higher scores values belonged to nitrogen-containing compounds (N-cnt), phenylpropanoids, terpenoids and lipids (Supplementary **Table S5**). As the treatment was the most discriminating factor in the AMOPLS-DA analysis, two study models were constructed that allow the separation between the different plant groups used in our study. One model compared ABA deficient plants (*flc*-) under any condition applied with WT plants under control conditions (named “*flc*- model”), and another model compared ABA deficient plants supplemented with exogenous ABA (*flc*+) with WT plants under control conditions (named “*flc*+ model”), being the last discussed in the next section.

Focusing in the “*flc*- model”, a Volcano analysis (Supplementary **Table S6**) identified 461 differentially accumulated metabolites (DAMs) in *flc*- grown under control, salinity, heat and salinity+heat from WT plants grown under control conditions. Using a Venn diagram, we identified independent and shared compounds between the studied treatments (**Fig. 3A**; (Bardou *et al*., 2014)). Only 28 DAMs overlapped among all environmental conditions, and only 26 DAMs overlapped for the stress treatments. Single stresses shared only 10 metabolites, while salinity and salinity+heat shared 35 DAMs, and heat and salinity+heat shared 86 DAMs. Thus, we can deduce that stress combination was primarily governed by the heat component at the metabolomic level. ABA deficiency induced significant changes at the metabolomic level in control conditions compared to WT and, as expected, these changes were more pronounced when *flc*- were subjected to stress conditions. Heat stress and stress combination had almost twice the number of metabolites.

**Fig. 3.**
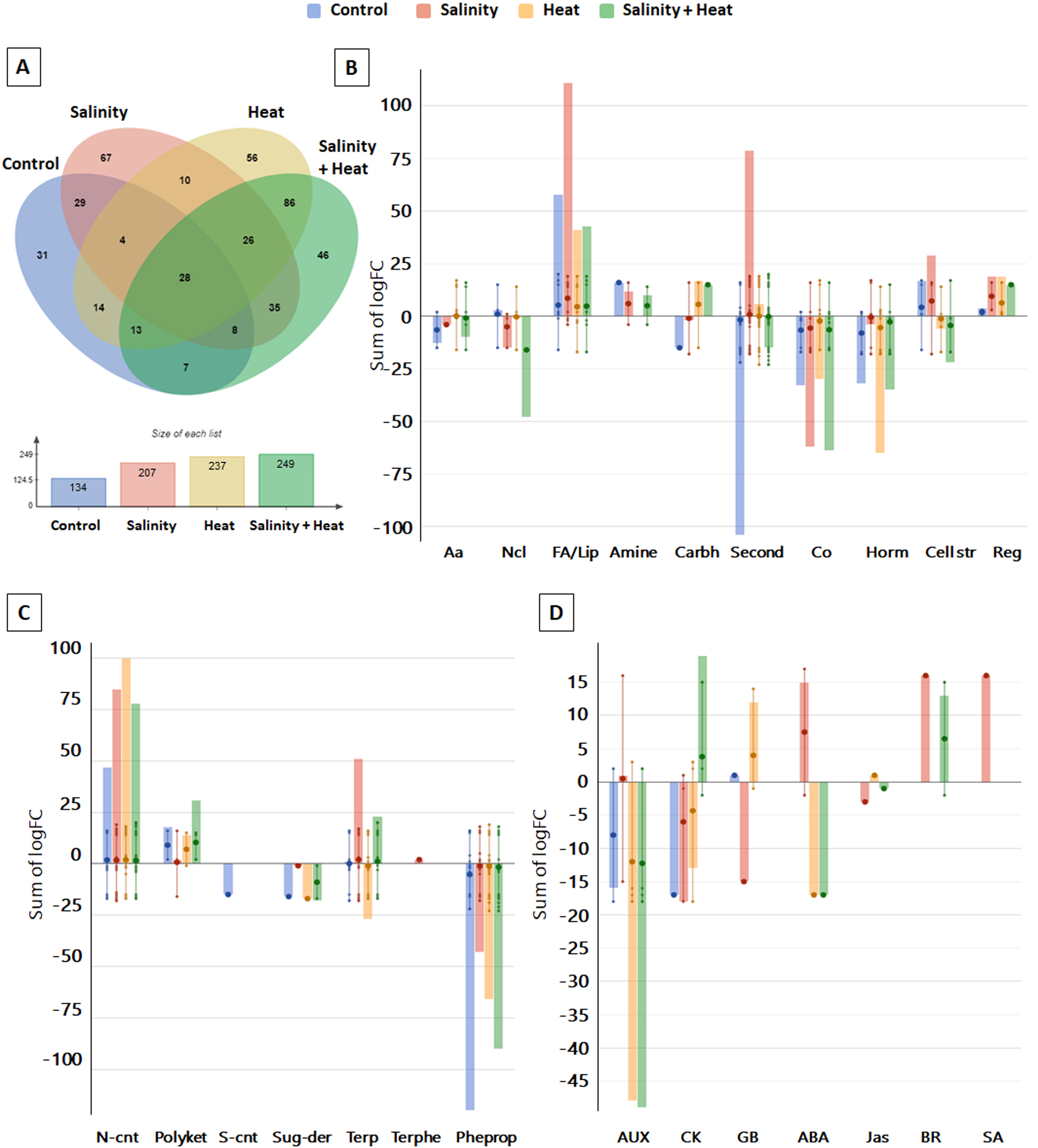
Differential accumulated metabolites (DAMs) in *flc-* mutants under control, salinity, heat and the combination of salinity+heat. and biosynthetic pathway analysis. (A) Venn diagrams of the overlap between DAMs in each treatment. (B) Biosynthetic pathway enrichment. (C) Secondary metabolite biosynthetic pathways analysis. (D) Hormone biosynthetic pathways enrichment. (B, C and D were performed using PlantCyc software). Data were normalized against WT control (p<0.05; FC>2). AA: amino acids; Ncl: nucleo; FA: fatty acids; Lip: lipids; Carbh: carbohydrates; Second: secondary metabolites; Co: cofactors; Horm: hormones; Cell str: cell structure; Reg: regulatory; N-cont: N-containing; Polyket: polyketides; S-cont: S-containing; Sug-der: sugar derivatives; Terp: terpenoids; Terphe: terpenophenolics; Pheprop: phenylpropanoids; AUX: auxins; CK: cytokinins; GB: gibberellins; ABA: abscisic acid; Jas: jasmonates; BR: brassinosteroids; SA: salicylic acid.

Continuing this analysis, the 461 DAMs identified by the Volcano analysis (Supplementary **Table S6**) were used for the biological interpretation of our results by conducting a pathway tool analysis using PlantCyc software (**Fig. 3B, C and D**). In ABA-deficient mutants (*flc*-) compounds related to amino acid, cofactors and hormone biosynthesis pathways were reduced, while compounds related to lipid and regulatory metabolite biosynthetic pathways were increased compared to WT under control condition (**Fig. 3B**). The results also showed a general increase in N-containing metabolites (N-cnt) and a general decrease in phenylpropanoids (Pheprop) in *flc*- plants. However, terpene (Terp) accumulation depended specifically on the environmental conditions to which *flc*- plants were subjected, increasing under salinity stress and stress combination and decreasing under heat stress (**Fig. 3C**). At the hormonal level, metabolites related to auxin (AUX) biosynthesis were significantly decreased in *flc*- plants grown under control conditions, heat stress and stress combination as compared to WT control. On the other hand, cytokinins (CK) exhibited different regulation, decreasing under control and single stress conditions and increasing under the stress combination (**Fig. 3D**). Overall our results indicated that levels of carbohydrates, amino acids, lipids, secondary metabolites and hormones are regulated by environmental conditions (Krasavina *et al*., 2014; Planchet and Limami, 2015; Liu *et al*., 2019a; Jan *et al*., 2021; Waadt *et al*., 2022).

As mentioned earlier, secondary metabolite biosynthesis at the transcriptomic and metabolomic levels was decreased in *flc*- under control conditions and under the combination of salinity+heat. Under single stress (salinity or heat), a strong downregulation of genes belonging to secondary metabolism was observed at the transcriptomic, but this was not reflected in a lower concentration of the compounds they regulate at metabolomic level. Furthermore, the upregulation of the transcription of genes related to the biosynthesis of phenylpropaoid compounds was not consistent with the decrease observed in the phenylpropanoid levels in *flc*- plants under single salt stress and stress combination. Various signaling pathways and multiple regulatory mechanisms, such as transcriptional, post-transcriptional, post-translational, epigenetic, phytohormonal signaling, and responses to biotic and abiotic stresses, collectively modulate phenylpropanoid metabolism. Zhang et al. (2020) showed that various microRNAs (miRNAs) and other small RNAs can target structural genes or regulators within the phenylpropanoid metabolism pathway, thus regulating the accumulation of some specific phenylpropanoid compounds (Zhang *et al*., 2020). Based on these results, we confirm that higher transcription does not always lead to a higher accumulation of metabolites in a particular pathway. This underscores the importance of studying the different -omics and their interconnection as a whole within systems biology.

#### Multi-omics study on *flc*- model and identification of the main key markers for each environmental condition

A total of 27 phenotypic parameters, 20 ions, 28522 transcripts, and 1515 metabolites were identified and quantified in the three plant groups (WT, *flc*-, *flc*+) grown under control, salinity, heat and salinity+heat conditions. Due to great number of transcripts and metabolites, DEGs and DAMs with a FC≥2 and p-value 0.05 were selected through a Volcano analysis (Supplementary **Table S2 and** Supplementary **Table S6**). DEGs, DAMs, ions and phenotypic parameters were subjected to a multi-omics analysis employing the MixOmic tool (Rohart *et al*., 2017). From these analyses, three models were designed, based on the plant group: *flc*-, *flc*+ and WT.

The resulting “*flc*- model” from the multi-omics analysis is presented in **Fig. 4**. Initially, the four datasets (phenomics, ionomics, transcriptomics and metabolomics) were individually plotted based on their scores on the first to components to evaluate their distribution before the multi-omics analysis **(Fig. 4A).** The sPLS-DA analysis (**Fig. 4A**) showed that “component 1” clearly separated control and salinity from heat and salinity+heat, while “component 2” discriminated control and heat from salinity and salinity+heat conditions. This separation was consistent with the HCA (**Fig. 4B**). When the four datasets were combined in the same analysis, Pearson correlation coefficients exceeded 0.9 for both components 1 and 2 (Supplementary **Fig. S4A and** Supplementary **S4B**), which validated our previous observations. Interestingly, when transcriptomic and metabolomic data were studied separately, a high number of DEGs and DAMs were common between heat and stress combination (**Fig. 2A and 3A**, respectively). However, when considering these two datasets along with ionomics and phenomics, as well as the influence of each parameter on a multi-omics analysis, the results showed that the general cellular response induced under salinity+heat stress was actually more similar to salinity than to heat (**Fig. 4B**). These results underscored the importance of combining the results from the different –omics analyses compared to individual –omics studies.

**Fig. 4.**
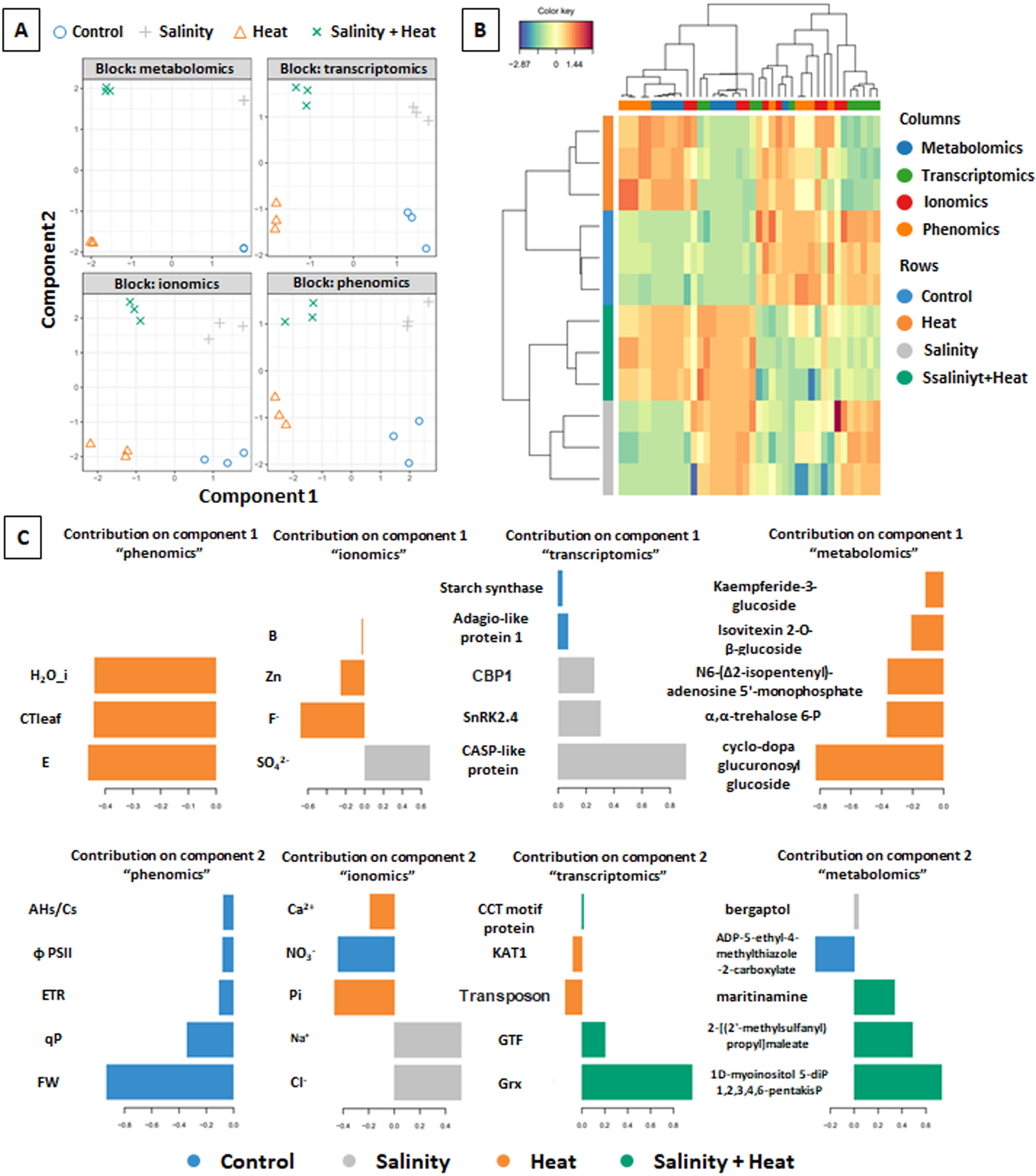
Multi-omics analysis in *flc*- mutant using MixOmics software. (A) sPLS-DA analysis in each –omic. (B) Heatmap correlation plot of most discriminant features selected from phenomic, ionomic, transcriptomic and metabolomic datasets. Omics are represented in columns and treatments are represented in rows. (C) Loading plot of each feature selected having the maximal discrimination ability on the first and second components in each -omics datasets. Color indicates the stress treatment applied. CBP1: CCG-binding protein1; SnRK2.4 sucrose non-fermenting related protein kinases 2.4; KAT1 K+ transporter: GTF: Glycosyltransferase; Grx: glutaredoxin.

Using mixOmics software facilitated the extraction of molecular markers from several large datasets in a specific genotype, statistically interconnecting them as a whole, and thus facilitating a biological interpretation of these markers. Consequently, molecular markers from each–omics dataset under a specific environmental condition were obtained based on their scores on the first two components for our “*flc*- model” (**Fig. 4C**).

For the phenomics, all molecular markers were associated with control conditions, with the most relevant ones being FW, NO_3_^-^, PSII (already identified in the individual phenomic and ionomic studies), ETR, qP and AHs/Cs. AHs/Cs takes into account CO_2_ assimilation, with respect to humidity and CO_2_ concentration of the leaf surface (Wolf *et al*., 2006), which it was positively correlated with the stomatal conductance and transpiration rate obtained for these *flc* mutants (**Fig. 1**). At the transcriptomic level, overexpression of starch synthase (SOLYC03G083090.4) and adagio-like protein (SOLYC01G005300.4) were the molecular markers obtained under control conditions. Adagio-like protein codes for a protein involved in photosynthesis (Wang *et al*., 2023) and, together with all the photosynthetic parameters mentioned, demonstrated a specific reprogramming of photosynthesis in *flc*-plants due to ABA deficiency. On the other hand, overexpression of a starch synthase could indicate that plants are investing in increasing starch levels (Thalmann and Santelia, 2017).

For salinity stress, several key markers were identified in *flc*- plants, including SO ^2-^, CCG-binding protein1 (CBP1, SOLYC01G108910), sucrose non-fermenting related protein kinases 2.4 (SnRK2.4, SOLYC02G090390), casparian strip membrane domanin proteins (CASP-like protein 4, SOLYC04G005555), along with the metabolite bergaptol. Si, SO_4_ ^2-^ and certain CASP-like proteins have been previously associated with salt stress tolerance in plants. Additionally, SnRK1α has been implicated in the ABA signalling pathway to improve tomato salt tolerance (Raziq *et al*., 2022). Interestingly, in ABA-deficient tomato plants (*notabilis*) an increase in the expression of SlSnRK2.4 was observed under soil flooding stress (De Ollas *et al*., 2021), suggesting its involvement in the ABA-independent salinity stress response.

For heat stress in *flc*- mutants, key markers included humidity (H20_i), leaf temperature (CTleaf), transpiration (E), Zn^2+^, F^-^, Ca^2+^ and Pi, along with overexpression of kat1 and increased levels of certain metabolites, such as kaempferide-3-glucoside, isovitexin 2-O-β-glucoside, trehalose 6-phosphate, adenosine 5’-monophosphate and cyclo-dopa glucuronosyl glucoside. Most of these markers have been previously associated with heat stress in plants (Wiebe and Poovaiah, 1973; Dias and Lidon, 2009; Pacak *et al*., 2016; Zheng *et al*., 2020; Laoué *et al*., 2022; Samineni *et al*., 2022; Wei *et al*., 2022). Notably, the K+ transporter (KAT1) is involved in K^+^ uptake in guard cells, regulating stomatal opening or closing and transpiration (Hosy *et al*., 2003). Its overexpression appears to be both ABA-dependent and ABA-independent (Mulet *et al*., 2023), appearing to have an ABA-independent overexpression in our *flc*- mutants. The metabolites identified, particularly phenylpropanoids generally play roles as antioxidants in the plant’s response to oxidative stress. However, the phenylpropanoids that have resulted from our analysis were in their glycosidated forms (kaempferide-3- glucoside, isovitexin 2-O-β-glucoside) and, therefore, may be not playing these antioxidant roles in our mutants (Le Roy *et al*., 2016).

Finally, for the stress combination, specific genetic and metabolic markers identified included the overexpression of CCT-motif protein (SOLYC03G083400.3), a glycosyltransferase (SOLYC04G074380. 4), and a glutaredoxin (SOLYC06G008760.1), and the increase of the compounds maritinamine (oxidoreductant compound), 2-(2’-methylsulfanyl)propyl]maleate and 1D-myo-inositol 5-diphosphate 1,2,3,4,6- pentakisphosphate. CCT motif protein has been associated with carotenoid pathway gene expression (Ye *et al*., 2015), while glutaredoxins has been shown to enhance plant tolerance to some type of stresses, such as salinity and heat (Li, 2014; Sousa *et al*., 2022). In our results, the glutaredoxin identified (SOLYC06G008760.1) was specifically regulated in *flc*- mutants subjected to stress combination, which might indicate its specific role under these conditions. Maritinamine accumulation, involved in oxido-reduction reactions, may contribute to detoxification of ROS in ABA-deficient mutant under stress combination (Kilgore *et al*., 2016). Maritinamine accumulation was also identified among the VIPs within the plant group x treatment interaction. Thus, it appears that maritinamine accumulation is specific in ABA-deficient plants under salinity+heat.

### Sufficient ABA levels in ABA-deficient mutant do not restore WT metabolism: flc+ model

#### Phenomics in *flc*+

From the first studies to date, it has been seen that the ABA application improves plant tolerance to different abiotic stresses (Itai *et al*., 1978; Rehman *et al*., 2021). In our studies, the exogenous application of ABA (*flc*+) indeed improved the phenotype and stress tolerance of *flc*- mutants, although did not fully restore the WT phenotype. This observation is consistent with previous studies that have shown the beneficial effects of ABA application on plants stress tolerance in *flacca* mutants (Imber and Tal, 1970; Li and Liu, 2021). Interestingly, the combination of salinity+heat resulted in highest biomass reduction in *flc*+ mutants (**Fig. 1A**), indicating a greater susceptibility to stress combination as compared to single stress. *Flc*+ plants also showed a significant increase in ABA levels respect to WT under single stress or stress combination suggesting that exogenous application of ABA led to elevated ABA level in these mutants. However, even with increased ABA levels, *flc*+ plants did not fully recover WT phenotype, indicating that other factors may contribute to the differences observed (**Fig. 1A**). Furthermore, other physiological parameters, including Ci/Ca under heat stress and H20_i and leaf temperature (CTleaf and SVTleaf) under control and salinity+heat combination, were significantly different between WT and *flc*+ plants (**Fig. 1A**; Supplementary **Table S1**), highlighting the complex interplay between ABA deficiency, ABA application, and environmental stresses in modulating plant physiology and stress responses.

#### Ionomics in *flc*+

At ionomic level, NO ^-^ and Pi concentrations were similar to those found in *flc*- under control conditions. The same observations were confirmed for Zn^2+^ under salinity and salinity+heat and SO_4_^2-^ under control, heat and salinity+heat conditions (**Fig. 1C**; Supplementary **Table S1**). Sagervanshi et al (2020) showed that the priming application of ABA to salt stress led to an increase in ions such as Zn^2+^, NO_3_ ^-^ and SO_4_ ^2-^ in *Vicia faba* plants (Sagervanshi *et al*., 2021). Our results are consistent with previous findings regarding Zn^2+^ and SO_4_ ^2-^ ions, establishing a relationship between the exogenous application of ABA and these ions. This effect also occurs under salinity+heat in *flc*+ plants. On the other side, in our *flc* tomato plants, differences in NO_3_ ^-^ levels between ABA application (*flc*+) and no application (*flc*-) under saline conditions were not observed. Possibly, these changes are due to the fact that our study was done on ABA-deficient tomato plants and Sagervanshi et al. (2020) on WT *Vicia faba* plants (Sagervanshi *et al*., 2021). Contrarily, ABA application increased Pi levels. Zhang et al. (2022) proved that ABA positively regulates Pi acquisition through ABI5 in the *Arabidopsis* response to phosphate deficiency (Zhang *et al*., 2022). Hence, NO_3_^-^, Pi, Zn^2+^and SO_4_ ^2-^ levels in tomato leaves seem to depend on ABA endogenous levels.

#### Transcriptomics in *flc*+

When studying the transcriptome response obtained from *flc*+ plants grown under control, salinity, heat and salinity+heat at total of 903, 2908, 6496 and 7423 DEGs, respectively, were obtained compared with WT plants grown under control condition (**Fig. 5A and** Supplementary **Table S7**). Romero et al. (2012) demonstrated that the application of ABA in sweet orange plant fruits from ABA-deficient mutant plants under dehydration condition did not produce a recovery at the transcriptomic level with respect to WT, supporting the results found in these experiments (Romero *et al*., 2012). This was observed with 903 DEGs compared to WT under control conditions and it was also noted in a previous work of our group (Pardo-Hernández *et al*., 2024). On the other hand, a certain number of DEGs were treatment-specific, obtaining 117, 567, 774 and 1426 DEGs specifically regulated under control, salinity, heat and stress combination respectively. In addition, single and stress combination treatments shared with control treatment a total of 384 DEGs, indicating that these genes were still *flc*+ genotype- dependent transcripts. It is also worth mentioning that only 100 DEGs were shared by the application of single stresses, 345 DEGs were shared by salinity and stress combination and that the highest number of DEGs (3787) were shared by heat and salinity+heat (**Fig. 5A**).

**Fig. 5.**
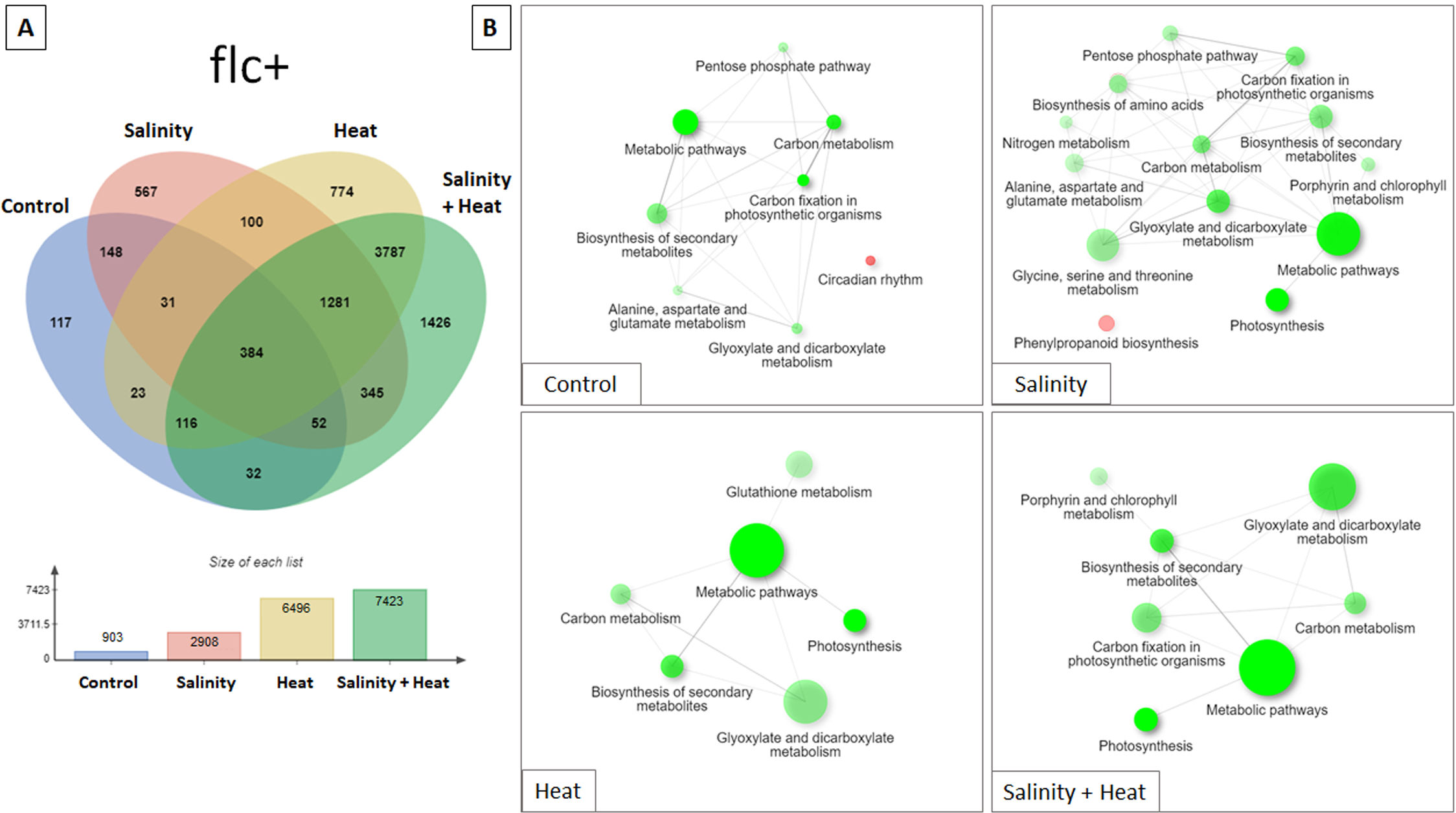
Differentially expressed genes (DEGs) in *flc*+ mutants under control, salinity, heat, and the combination of salinity+heat. (A) Venn diagrams illustrating the overlap between DEGs in each stress treatment. DEGs were extracted through normalization against WT control (p < 0.05; FC > 2). (B) KEGG pathway enrichment analysis of specific DEGs obtained under control, salinity, heat, and salinity+heat treatments in *flc*+ plants. The size of the circles corresponds to the number of genes associated with each pathway, and the intensity of the color reflects the significance level of the pathway (P < 0.05), with red representing upregulated DEGs and green representing downregulated DEGs.

KEGG enrichment pathway analysis on *flc*+ under the four environmental conditions used (**Fig. 5B**; Supplementary **Fig. S1**) showed a downregulation of the biosynthesis of secondary metabolism, carbon metabolism and glyoxylate and dicarboxylate metabolism. Additionally, *flc*+ mutants grown under single and stress combination showed a downregulation of the photosynthesis pathway. However, many KEGG pathways were generally downregulated under the different stress treatments applied. Thus, *flc*+ mutants grown under control or under salinity showed a specific downregulation of pentose phosphate pathway and alanine, aspartate and glutamate metabolism. On the other hand, downregulation of carbon fixation was observed in *flc*+ mutants grown under control, salinity and salinity+heat conditions. Finally, a specific downregulation of porphyrin and chlorophyll metabolism was observed when these plants were grown under salinity and salinity+heat combination. Controversely, several KEGG pathways were specifically enriched depending on the stress applied. For example, (i) circadian rhythm-related genes were specifically upregulated in *flc*+ grown under control; (ii) amino acids biosynthesis, nitrogen metabolism, and glycine, serine and threonine metabolism were specifically downregulated under salinity, whereas phenylpropanoid biosynthesis was upregulated; (iii) and, glutathione metabolism was specifically downregulated under heat.

#### Metabolomics in *flc*+

As mentioned in section 1.4, the metabolomics analysis annotated 1515 compounds (Supplementary **Table S3**) and both, unsupervised (HCA, Supplementary **Fig. S2**) and supervised (AMOPLS-DA, Supplementary **Table S4;** Supplementary **Fig. S3**) studies considering the three plant groups (WT, *flc*- and *flc*+) and the four treatments (control, salinity, heat, salinity+heat) were performed as explained in the “*flc*- model”. Concerning “*flc*+ model”, the principal components (tp5 and tp6) that identified for the interaction between treatment and genotype (plant group x treatment) showed that *flc*+ plants were significantly different from WT under the four treatments (Supplementary **Fig. S3C**).

A Volcano analysis (Supplementary **Table S8**) identified 458 DAMs in *flc*+ that significantly differed from those found in WT plants under control conditions. We were able to identify the specific and common compounds between the treatments applied to *flc*+ using a Venn diagram (**Fig. 6A)**. Only 28 DAMs overlap for all the environmental conditions applied, and only 35 metabolites overlap for the stress treatments. Single stresses shared only 13 DAMs, while salinity and salinity+heat shared 33 DAMs, and heat and salinity+heat shared 103 DAMs. Similarly, to the observed in the previous “*flc*- model”, stress combination was governed by the heat component at the metabolomic level.

**Fig. 6.**
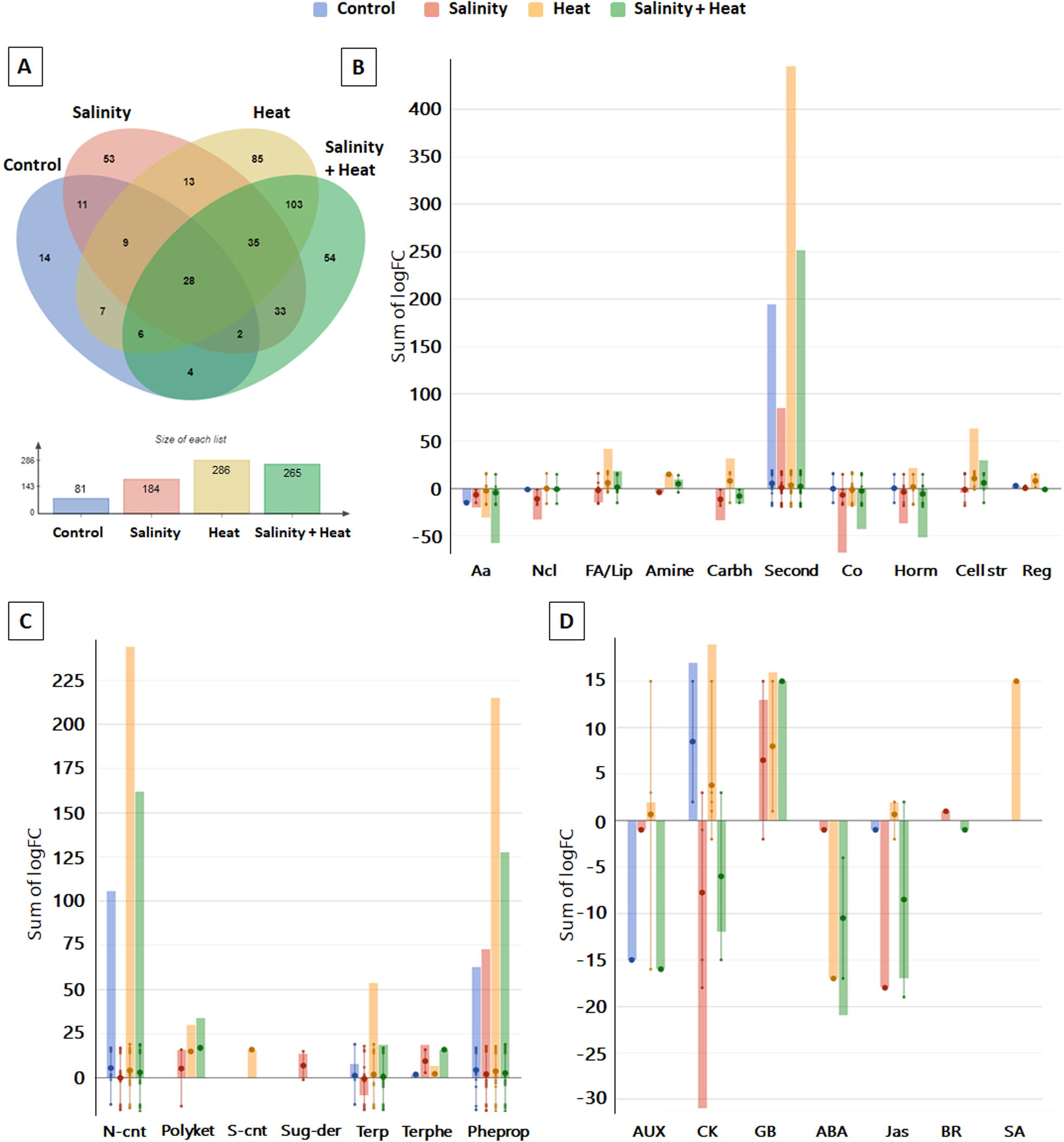
Differential accumulated metabolites (DAMs) and biosynthetic pathway analysis in *flc*+ mutants under control, salinity, heat, and the combination of salinity+heat. DAMs were extracted by normalization against WT control (p < 0.05; FC > 2). (A) Venn diagrams illustrating the overlap between DAMs in each treatment. (B) Biosynthetic pathway analysis. (C) Secondary metabolites biosynthetic pathways analysis. (D) Hormone biosynthetic pathways analysis. (B, C, and D were analyzed using PlantCyc software). Abbreviations: AA: amino acids; Ncl: nucleotides; FA: fatty acids; Lip: lipids; Carbh: carbohydrates; Second: secondary metabolites; Co: cofactors; Horm: hormones; Cell str: cell structure; Reg: regulatory; N-cont: N-containing; Polyket: polyketides; S-cont: S-containing; Sug-der: sugar derivatives; Terp: terpenoids; Terphe: terpenophenolics; Pheprop: phenylpropanoids; AUX: auxins; CK: cytokinins; GB: gibberellins; ABA: abscisic acid; Jas: jasmonates; BR: brassinosteroids; SA: salicylic acid.

Subsequently, the 458 DAMs identified by the Volcano plot analysis (Supplementary **Table S8**) were used for pathway analysis. An increase in secondary metabolism was observed, especially in those plants grown under heat stress (**Fig. 6B**). Upon inspecting the DAMs belonging to the secondary metabolism, a general increase in phenylpropanoid-related metabolites was observed, especially in *flc*+ plants grown under heat stress. In addition, a significant accumulation of N-cont was observed, as we previously observed in the “*flc*- model”. However, in *flc*+ salinity stress did not induce these differences observed in the “*flc*- model” with respect to WT plants. Interestingly, and in contrast to the “*flc*- model”, Terp were less abundant in *flc*+ plants under salinity stress (**Fig. 6C)**. At the hormonal level, metabolites related to auxin and cytokinin biosynthesis pathways were significantly decreased in *flc*+ plants under control and salinity+heat combination and, in the case of cytokinins, a decrease of these metabolites were observed under single stresses (heat or salinity) (**Fig. 6D)**. Watanabe et al. (2018) demonstrated that *aba3* mutants of *Arabidopsis*, an ABA-deficient mutant homolog to *flc* in tomato, showed a decrease in anthocyanins that was not recovered by ABA application (Watanabe *et al*., 2018). Thus, our results showed that in our experiments, ABA endogenous concentration was not related with the plant response under control and heat stress conditions.

#### Multi-omics analysis in *flc*+ model and identification of the main key markers for each environmental condition

As in the *flc*- model, DEGs and DAMs with a FC≥2 and p-value 0.05 identified in flc+ model were selected through the Volcano analysis (Supplementary **Table S7 and** Supplementary **Table S8**) and, together with the phenomics and ionomics, were all submitted to the multi-omics analysis. The four datasets (phenomics, ionomics, transcriptomics and metabolomics) were initially plotted individually according to their scores on the first two component through sPLS-DA analysis (**Fig. 7A**) (Rohart *et al*., 2017). Additionally, Pearson correlation coefficients obtained from the analysis of all - omics datasets together exceeded 0.9 for both components (Supplementary **Fig. S4C and** Supplementary **S4D**), as it was previously described for the “*flc*- model”. Also, the same observations made previously for *flc*- model respect to the treatment separations through an HCA analysis were observed in *flc*+ model, (**Fig. 7A**; **Fig. 7B**) showing control and heat, on one side, and salinity and salinity+heat, on the other side. Here again, the study of the different –omics as a whole is shown as a fundamental key to ensuring the effect of a given stress condition to the entire plant metabolism (**Fig. 6A**; **Fig. 6A**; **Fig. 7C**; Supplementary **Table S11**).

**Fig. 7.**
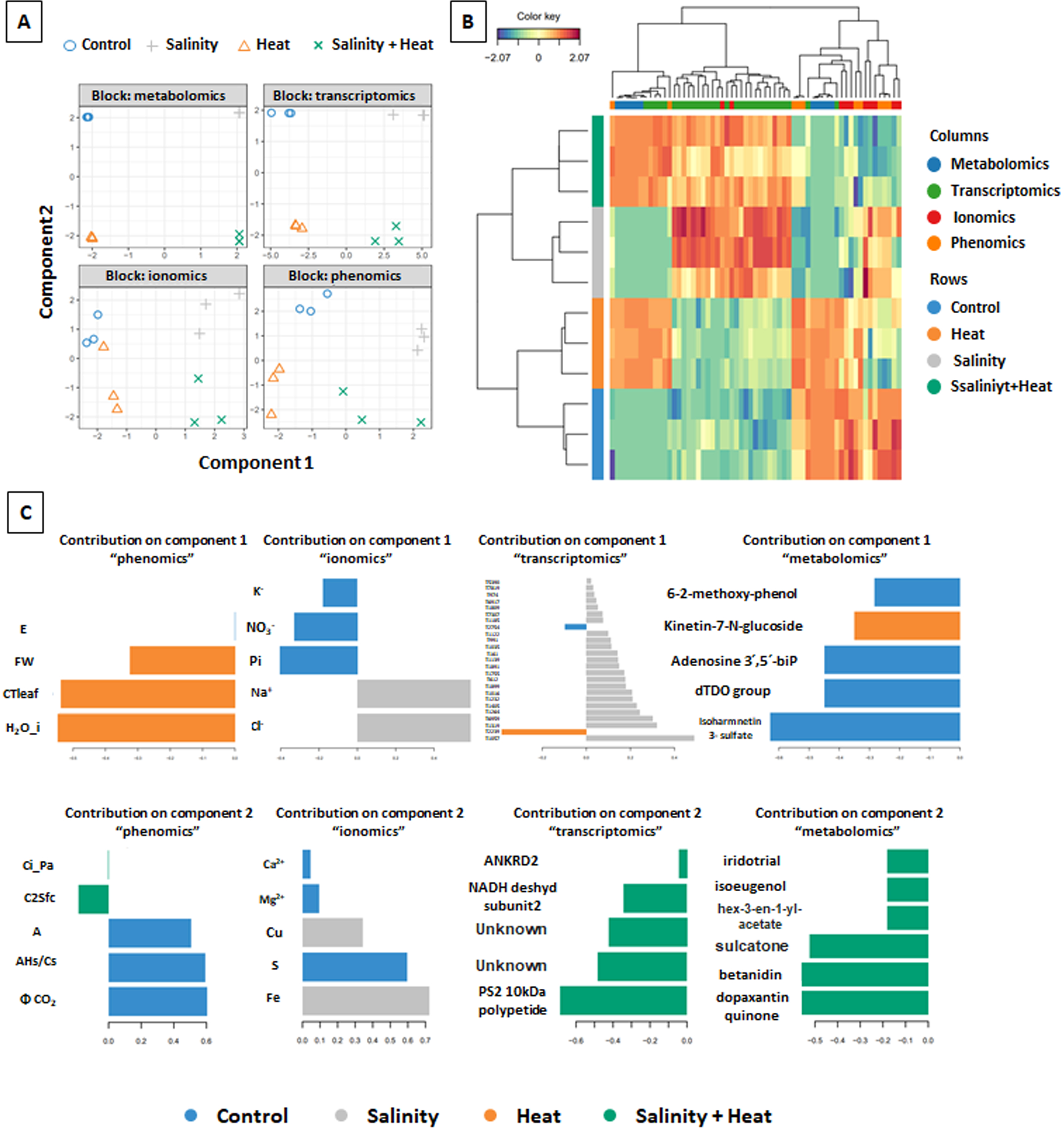
Multi-omics analysis in *flc*+ mutant using MixOmics software. (A) sPLS-DA analysis in each –omic. (B) Heatmap correlation plot of most discriminant features selected from phenomic, ionomic, transcriptomic and metabolomic datasets. Omics are represented in columns and treatments are represented in rows. (C) Loading plot of each feature selected having the maximal discrimination ability on the first and second components in each -omics datasets. Color indicates the stress treatment applied.

Under control conditions, *flc*+ plants were associated to two key photosynthetic markers, CO_2_, *A*, K^+^, NO_3_^-^, Pi, Ca^2+^, Mg^2+^, S. In addition, a peptidylprolyl isomerase (SOLYC09G008650.3) was also revealed as an associated marker at a transcriptomic level, which could be involved in protein folding after stress-induced damage (Bizouerne *et al*., 2023). Additionally, metabolites such as isorhamnetin 3-sulphate (persicarin), the first sulfated flavonoid found in plants (Harborne, 1975), were identified as key markers in *flc*+ under control conditions. The function of flavonoid sulfation is still unclear, but it has been shown that sulfated flavonoids may be involved in molecular recognition and detoxification of some signalling pathways (Teles *et al*., 2018).

The markers identified under salinity stress in *flc*+ mutants were at the ionic level Na^+^, Cl^-^, Cu^2+^ and Fe^2+^. It must be noted that the last two ions (Cu^2+^ and Fe^2+^) were not previously identified as putative markers when the ionomics were analysed as a separate dataset. Cu^2+^ and Fe^2+^ are essential cofactors in plant antioxidant enzymes, thus, protecting against oxidative stress (Bhaduri and Fulekar, 2012; Verma, 2016). Additionally, two transcription factors (bHLH transcription factor and MYB48), an aquaporin, a nitrogen transporter and several genes related to stress response, such as the ABA and environmental stress-inducible protein TAS14 (Supplementary **Table S11**) were identified as key markers in *flc*+ under salinity. Both, changes in ions and in the transcription level of genes related to ion transport, have usually been related to salt stress (Lopez-Delacalle *et al*., 2020; Amin *et al*., 2021) which help to confirm the results obtained.

The heat stress markers identified in *flc*+ were an increase in leaf FW, leaf temperature and cellular water content (as expected and due an increase in the transpiration rate to reduce leaf temperature under these conditions, which might induce a higher CO_2_ assimilation rate and higher growth). Additionally, an increase in the concentration of the metabolite kinetin-7-N-glucoside, which has previously been associated with tolerance to high temperature stress in plants (Itai *et al*., 1978; Osman *et al*., 2016; Mei *et al*., 2023) was observed. However, we are not sure of a given role to this marker under heat stress in our *flc*+ plants, since glycolsylated forms of cytokinins has been recently related to their inactivation (Li *et al*., 2019; Hluska *et al*., 2021). Thus, more research is needed in the role of this metabolite in heat stress tolerance.

Finally, the stress combination again showed a specific regulation with the identification of some specific markers that were not found in these plants when these stresses were applied individually. Both intercellular (*Ci*) and leaf surface (C2Sfc) CO_2_ levels were significantly higher under these conditions and an upregulation of photosystem II 10kDa polypeptide, chloroplastic (SOLYC12G017250.2) was observed, indicating a possible positive and specific regulation of photosynthesis. Upregulation of photosystem II 10kDa polypeptide has been associated with salt stress tolerance in tomato roots (Li *et al*., 2022). However, in our case, this protein was identified as a marker for salinity+heat stress combination in tomato leaves, possibly due to exogenous ABA application in our *flc* mutants or due to a dependence on the tomato plant organ studied. Also, the transcription of NADH dehydrogenase subunit 2 (SOLYC03G063500.1), ankyrin repeat domain-containing protein 2 (SOLYC10G024330.3) and two unknown proteins (SOLYC08G036540.1 and SOLYC07G021155.1) were identified as markers for this stress combination. Additionally, two compounds related to betalain biosynthesis (betanidin and dopaxanthin quinone) were identified as metabolic markers in *flc*+ under stress combination, and, curiously, these two markers were previously identified as VIPs of the plant group x treatment interaction in the metabolomic study (Supplementary **Table S5**). In addition, isoeugenol was also identified as a VIP for the treatment group in the independent metabolomics study (Supplementary **Table S5**) and, in our multi-omics study appeared again as a marker for stress combination. However, these betalains and isoeugenol have been related to the response to abiotic stresses but never to the combination of salinity + high temperature (Sokolova *et al*., 2022; Yeshi *et al*., 2022; Zhao *et al*., 2022), which highlight the importance of our studies.

#### Validation of flc- and flc+ molecular markers through the “Wt model”

After conducting multi-omics analyses in both *flc*- and *flc*+ models, we proceeded to perform similar analyses in WT plants under single or combined stresses with the aim of validating the specificity or the general association of these markers with the stress applied or the genotype studied. Thus, a similar study than those described previously by *flc*- and *flc*+ was performed with WT plants (Supplementary **Table S9;** Supplementary **Table S10;** Supplementary **Fig. S4E;** Supplementary **Fig. S4F;** Supplementary **Fig. S5A**). HCA showed two different groups: control and salinity, from one side, and heat and salinity+heat (Supplementary **Fig. S5B**). Li et al. (2023) demonstrated that heat played a higher contribution in transcriptomic and metabolomics levels when jointly applying heat and salinity stresses to wild tomato plants (Li *et al*., 2023). Previous results from our research group in commercial tomato lines also demonstrated these findings (Lopez-Delacalle *et al*., 2020, 2021). However, in the *flc*- and *flc*+ models the stress combination was more associated with salinity stress, therefore, our results showed a putative genotype-specificity on the abiotic stress tolerance mechanisms (**Fig. 4B and 7B**). The most relevant key markers associated with physiological parameters, ions, transcripts and metabolites identified in the “WT model” are reported in Supplementary **Fig. S5C**.

By comparing the markers obtained in the WT model (**Fig. 4C**; **Fig. 7C**; Supplementary **Fig. S5C;** Supplementary **Table S11**) with those obtained previously for *flc*- and *flc*+ we found several common markers (**Fig. 8)**. The markers specific to control conditions in at least two of the three plant groups were AHs/Cs, NO_3_^-^, PSII, ETR, qP, A and CO_2_, the last two being ABA-dependent because they were only present in WT and *flc*+ plants. The markers common to salinity stress were only Na^+^ and Cl^-^, which is logical considering that these ions were applied into the growing nutrient solution in this treatment. The markers obtained for heat stress were CTleaf, H_2_O_i and F^-^. Finally, Ci_Pa, C2Sfc, iridotrial, isoeugenol and (3Z)-hex-3-en-1-yl acetate (H-EN-YL-AC) were stress combination and ABA-dependent markers, as all of them were markers found in both WT and *flc*+ plants under this treatment.

**Fig. 8.**
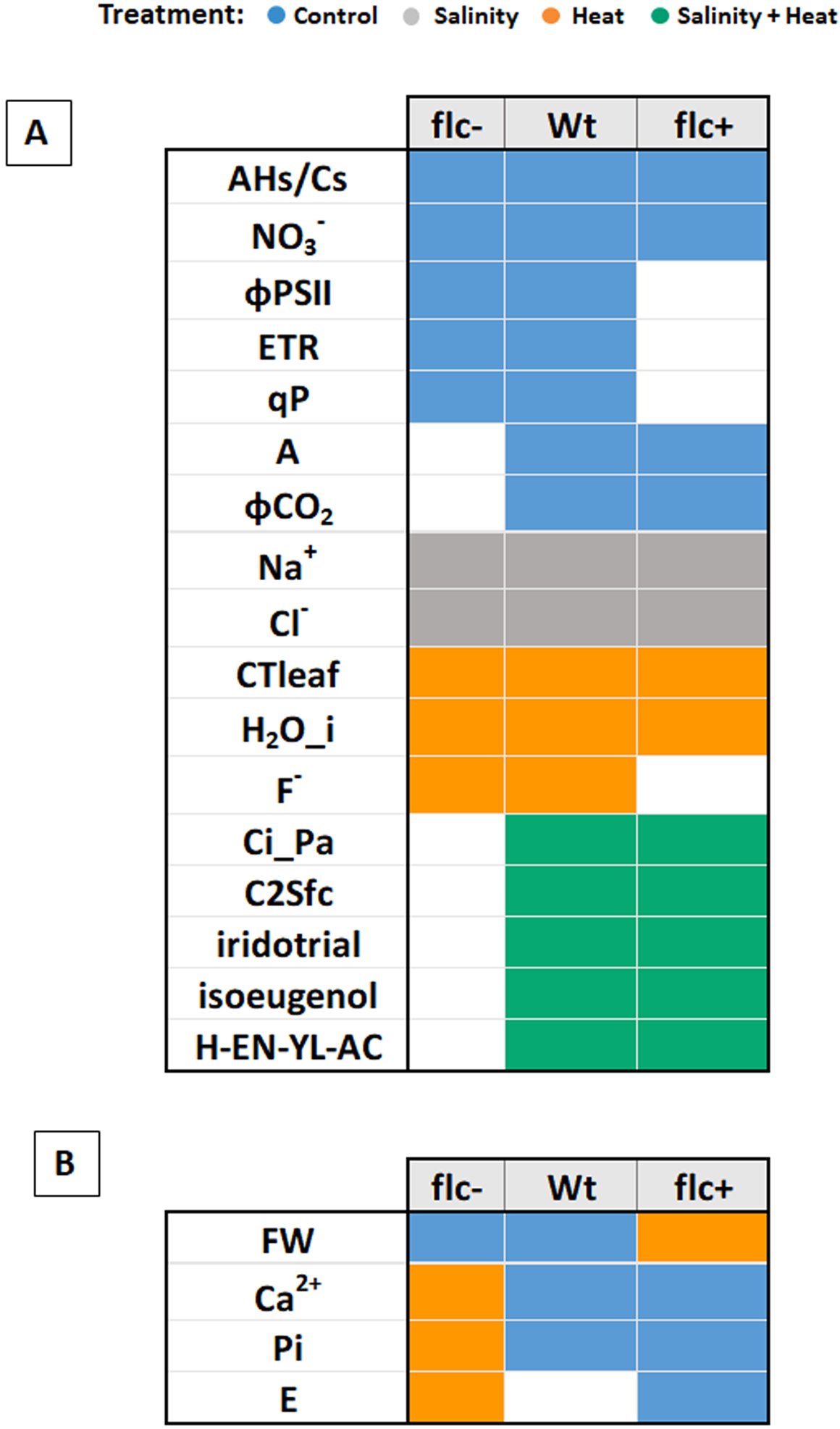
Multi-omics markers common in at least two of the three plant groups (WT, *flc*- or *flc*+). (A) Multi-omics markers that share the same treatment among different plant groups. (B) Multi-omics markers that do not share the same treatment among different plant groups.

Lastly, two physiological parameters (FW and E) and two ions (Ca^2+^ and Pi) were common markers in at least two of the plant groups, but, however, not under the same treatment. Ca^2+^ and Pi were markers in control conditions in WT and *flc*+ plants and markers of heat stress in *flc*- plants. The FW was a marker in *flc*- and WT plants under control conditions and in *flc*+ plants under heat stress. On the other hand, transpiration (E) was characteristic in *flc*- and *flc*+ plants, being a marker for heat and control conditions, respectively.

## Conclusions

ABA-deficient plants show significant changes in biomass, photosynthesis, ions, transcripts, and metabolites compared to wild-type plants under control conditions, salinity, heat, and salinity+heat. Our data demonstrate that some of these responses are independent of the optimal ABA endogenous concentration, thus being dependent on the applied environmental condition. On the other hand, exogenous application of ABA in ABA-deficient plants did not lead to plant recovery, neither at the phenotypic nor at the total metabolic level, but it does result in cellular reprogramming with changes in specific markers due to this exogenous ABA.

Plant response to environmental stresses is dependent on the stress applied, with a specific response to the application of single or combined stresses. Under field conditions, plants are subjected to various stresses simultaneously, and their response is not the sum of responses to individual stresses, but rather a specific response to that combination of stresses. Additionally, our multi-omics analyses have shown that each applied environmental condition presents specific markers of plant response. Markers observed in plants exposed to single stress (salinity and heat) differed from those observed in the combination of both stresses. Furthermore, here we have demonstrated that studying the different omics together enables the identification of certain specific markers for each stress condition that were not detectable in the study of each omics individually. Our results offer a new route for identifying specific metabolic markers in plant response to single stresses, and more importantly, to combined stresses, with implications for metabolic pathways, ion absorption, and physiological responses crucial for plant tolerance to climate change.

## Supplementary data

**Fig. S1**. Hierarchy clustering analysis (HCA) for transcriptomic analysis.

**Fig. S2.** Hierarchy clustering analysis (HCA) for metabolomics analysis.

**Fig. S3.** The AMOPLS scores plots corresponding to the predictive components (Tp) found in the metabolomic analysis associated with three factors: (A) Plant group (B) Treatment and (C) Plant group x treatment.

**Fig. S4.** Diagnostic sPLS of multi-omics analysis (A) component 1 in *flc*- model (B) component 2 in *flc*- model (C) component 1 in *flc*+ model (D) component 2 in *flc*+ model (E) component 1 in WT model (F) component 2 in WT model

**Fig. S5.** Multi-omics analysis in WT plants using MixOmics software. (A) sPLS-DA analysis in each –omic. (B) Heatmap correlation plot of most discriminant features selected from phenomic, ionomic, transcriptomic and metabolomic datasets. Omics are represented in columns and treatments are represented in rows. (C) Loading plot of each feature selected having the maximal discrimination ability on the first and second components in each -omics datasets. Color indicates the stress treatment applied.

**Table S1.** Phenomic and ionomic raw data

**Table S2.** DEGs of *flc*- model

**Table S3.** Metabolomic raw data

**Table S4.** Relative variability and block contributions of the AMOPLS analysis of metabolomic data acquired from tomato wild-type (WT) and *flacca* mutants without (*flc*-) and with ABA application (*flc*+) under control (C), salinity (S), heat (H), and combination of salinity and heat (S + H). RSS: Relative Sum of Squares; RSR: Residual Structure Ratio; Tp: predictive components; To: orthogonal component.

**Table S5.** VIPs of metabolomics analysis

**Table S6.** DAMs of *flc*- model

**Table S7.** DEGs of *flc*+ model

**Table S8.** DAMs of *flc*+ model

**Table S9.** DEGs of WT model

**Table S10.** DAMs of WT model

**Table S11.** Transcripts and metabolites IDs and their corresponding identification nomenclature for the key markers obtained in the multi-omics in *flc*+ and WT models

**File S1.** References on the detailed description of the UHPLC/QTOF-HRMS approach applied on zucchini exocarps samples.

## Acknowledgments

We sincerely acknowledge Mario G. Fon for proofreading the manuscript.

## Author contributions

MP-H conducted the experiments, performed phenomics, ionomics, transcriptomics, metabolomics, multi-omics, and statistical analyses, wrote the manuscript, and designed the figures and tables. PP-G conducted the metabolomics and multi-omics analyses, contributed to the writing, and designed some figures. LL contributed to the writing, editing, data analysis, and literature review. RMR conceived the project, supervised and analyzed phenomics, ionomics, and RNA-seq data, contributed to designing the final figures, corrected and supervised the writing and editing of the manuscript. All authors contributed to the article and approved the submitted version.

## Conflict of interest

All authors declare no commercial, industrial links or affiliations.

## Funding

This research was supported by the Ministry of Economy and Competitiveness from Spain (MCIU/AEI/FEDER, UE, Grant No. PGC2018-09573- B-100) to RMR; by the Ministry of Science and Innovation of Spain (Ph.D. Fellowship Grant No. FPU20/03051 and Mobility Grant No. EST23/00700) to MP-H. The authors thank the “Romeo ed Enrica Invernizzi Foundation”, Milan (Italy) for the kind support to the metabolomics facility at Università Cattolica del Sacro Cuore.

## Data availability

The phenomic and ionomic raw data can be found in the Supplemental Information provided and available at Plant Communications Online System. The RNA-seq reads are available National Center for Biotechnology Information (NCBI) database under the Sequence Read Archive (SRA, https://submit.ncbi.nlm.nih.gov/subs/sra/), under the BioProject identification number PRJNA947059 (Submission ID: SUB12991832; direct link to datasets: https://www.ncbi.nlm.nih.gov/sra/PRJNA947059). Metabolomics data have been deposited to the EMBL-EBI MetaboLights database (DOI: 10.1093/nar/gkad1045, PMID:37971328) with the identifier MTBLS9686.

## Notes

### Competing Interest Statement

The authors have declared no competing interest.

